# Transcriptomics predicts Artificial Light at Night’s (ALAN) impact on fitness: nightly illumination alters gene expression pattern and negatively affects fitness components in the midge *Chironomus riparius* (Diptera:Chironomidae)

**DOI:** 10.1101/2024.11.13.623161

**Authors:** Linda Eberhardt, Halina Binde Doria, Burak Bulut, Barbara Feldmeyer, Markus Pfenninger

## Abstract

The emission of artificial light at night (ALAN) is rapidly increasing worldwide. Yet, evidence for its detrimental effects on various species is accumulating. While the effects of ALAN on phenotypic traits have been widely investigated, effects on the molecular level are less well understood. Here we aimed to integrate the effects of ALAN at the transcriptomic and the phenotypic level. We tested these effects on *Chironomus riparius*, a multivoltine, holometabolous midge with high ecological relevance for which genomic resources are available. We performed life-cycle experiments in which we exposed midges to constant light and control conditions for one generation. We observed higher EmT50 and reduced fertility under ALAN. From the observed decline in population size due to the reduced fertility, we predicted the population size to decline to 1% after 200 days. The transcriptomic analysis revealed expression changes of genes related to circadian rhythmicity, moulting, catabolism and oxidative stress. From the transcriptomic analysis we hypothesised that under ALAN, oxidative stress is increased, and that moulting begins earlier. We were able to confirm both hypotheses in two posthoc experiments, showing that transcriptomics are a powerful tool in predicting physiological outcomes before they are even observable.

## 1. Introduction

Most organisms have some of their life-history traits synchronised with daily or seasonal photoperiodic cues (McLay *et al.,* 2017). The periodic cycling of light and dark drives feedback loops of the endogenous circadian clock (Abe *et al.,* 2022; Carmakian & Boivin, 2009). The rhythmically oscillating transcription of circadian clock genes then regulates the expression of traits such as pupation, eclosion, mating or metabolism (Franco *et al.,* 2017, in Brady et al., 2021, Feng & Lazar, 2012, Tomioka & Matsumoto, 2010, Saunders, 2002). Changes in this delicately balanced system are therefore likely to trigger changes in these traits, starting from changes in gene expression to impacts on population fitness, which may eventually lead to population demographic effects.

One of the many influences of human civilisation on the natural environment is local nightly illumination by anthropogenic light sources which disrupts natural dark periods (Owens & Lewis, 2018). In scientific literature, this anthropogenic nightly illumination is commonly referred to as artificial light at night (ALAN). Adverse, albeit variable across taxa, effects of the disruption of photoperiodic cues by ALAN have been confirmed in vertebrates (Robert *et al.,* 2015, in McLay *et al.,* 2017) and invertebrates, including moths (van Geffen *et al*., 2014), fruit flies (Thakurdas *et al.,* 2009), crickets (Durrant *et al.,* 2015) and ants (Lone & Sharma, 2008). Effects on these organisms’ fitness components included juvenile development (van Geffen *et al*. 2014), timing of eclosion (Thakurdas *et al.,* 2009) and fecundity (McLay *et al.,* 2017) with a nightly brightness increase as low as 10 Lux. As this extensive listing shows, potential reactions to ALAN are so diverse, that negative impacts on many traits may be easily missed, because they were not measured in the field or the experiment. In the current study, we tried to minimise this risk by exploiting the fact that most changes in higher level traits such as fitness components have their initial cause in gene expression changes By quantifying changes in gene expression patterns, we infer potentially affected traits and test the arising hypotheses post hoc with respective experiments. In other words, we use transcriptome analysis to predict and test adverse fitness effects.

*Chironomus riparius* (Meigen 1803) (Diptera: Chironomidae) is a widespread, multivoltine, holometabolous insect that spends all four larval stages and the pupal stage in the aquatic environment before emerging as the adult imago (Armitage *et al.,* 1995). *C. riparius* is an ecologically highly relevant species as many taxa in water, land and air rely on these midges as a food source (Armitage *et al.,* 1995). The short life cycle of around 30 days (Armitage *et al.,* 1995) makes it an ideal study subject for eco-evolutionary experiments (Doria et al., 2022, Waldvogel & Pfenninger, 2021, Pfenninger & Foucault, 2020, Foucault et al., 2019, Foucault et al., 2018), moreover, since comprehensive genomic resources are available (Schmidt et al., 2020). Given its ubiquitous appearance, the species also occurs in areas affected by ALAN. It has been shown that photoperiod affects fitness and developmental speed in *C. riparius* and that there is likely a genetic basis for seasonal adaptive response to changes in photoperiod (Doria et al., 2021). However, the effects of ALAN on *C. riparius* at various trait levels have not been studied thus far. Therefore, the aim of this study was to examine whether the presence of ALAN at a level of 100 Lux affects *C. riparius’* gene expression and fitness components; the most basal and the most highly integrated organismic phenotypic trait levels.

## 2. Material & Methods

### 2.1 Experimental design

The *C. riparius* population used in the study originated from a native population (Hasselbach, Hessen, Germany; 50.167562°N, 9.083542°E) and has been maintained for several generations as a large in-house laboratory culture that was regularly replenished from the field population. This population has been previously used in several ecotoxicology and eco-evolutionary studies (e.g. Rigano et al. 2024, Carrasco-Navarro et al., 2021, Doria and Pfenninger, 2021, Pfenninger and Foucault, 2020).

Life-cycle tests according to OECD guideline 233 (OECD, 2010) were employed to investigate whether ALAN affects development and reproduction in chironomids. Five freshly laid egg ropes from different parents were drawn from the culture, put into 6-well plates in 3 mL per well of medium, consisting of deionised water adjusted to a conductivity of 550 µS/cm with aquarium sea salt (e.g., TropicMarin®), and a basic pH around 8 (Foucault et al., 2019), and let to hatch. The fully hatched egg ropes were chosen to be tested under two different conditions in an optimal photoperiod of 16:8 light:dark. All treatments had an identical 16 hours daytime (1000 lux equivalent to an overcast day) followed by 8 hours night time of either natural darkness (0 lux) or (100 lux) ALAN treatment, corresponding to bright urban night lighting.

Tests were performed with fifteen replicates for each treatment, meaning that a glass bowl per replicate (Ø 20 × 10 cm) was filled with a 1.5 cm sediment layer of washed, pH neutral, commercial sand and 1.250 L of medium (Foucault et al., 2019). Experiments were initiated by adding 30 first instar larvae to each bowl. The larvae in each treatment group originated from the same egg clutch to exclude family effects (Foucault et al., 2019). Test vessels were kept at 20 °C, 60% humidity and were constantly aerated. Water evaporation in the test vessels was compensated by adding demineralised water. Conductivity was maintained at 550 - 650 µS/cm with a pH around 8. Th chironomids were fed with finely ground fish food (e.g., Tetramin® Flakes) according to an age-dependend feeding schedule (Supplementary Material S2).

### 2.2 RNA sequencing and gene expression analysis

To investigate the underlying effects of ALAN at the gene transcription level, we performed a total RNA sequencing analysis on five of the fifteen replicates per treatment. Six to eight L4 larvae from each replicate were collected and pooled together approximately two days before expected emergence. The pools were immediately homogenized in TRIzol (Invitrogen, ThermoFisher Scientific, Waltham MA, USA) with a pestle and stored at −80°C. RNA extraction of five pools per replicate was performed with Direct-zol RNA Miniprep (Zymo Research). RNA samples were stored at −80°C after extraction. Single-stranded messenger RNAs (mRNAs) enrichment and conversion to complementary DNA (cDNA), library preparation and sequencing of 150 bp paired end reads using Illumina NovaSeq platform were performed by Novogene Europe – UK.

The quality of the reads was assessed using Fastqc (https://www.bioinformatics.babraham.ac.uk/projects/fastqc/). Subsequent trimming of adapters was performed with Trimmomatic v0.3.9 (Bolger *et al.,* 2014) and the following settings: ILLUMINACLIP: adapter all.fa:2:30:10:8), HADCROP:10, SLIDINGWINDOW:4:20 and MINLEN:50. The quality of the trimmed reads was assessed again with Fastqc. 7.6% of reads were discarded in the trimming process in both groups.

Filtered reads were mapped against the *C. riparius* (Laufer) draft genome v.3 (Schmidt *et al.,* 2020) using Hisat2 v.2.1.0 (Kim *et al.,* 2019). The gff files were converted into gtf format using gffread v.0.12.7 (Pertea & Pertea, 2020). A putative splice sites file was created using Python and the toolkit hisat 2 with the command extract_splice_sites.py. Subsequently, mapping was performed using Hisat2 with the generated splice-site file using the mode *–known-splicesite-infile.* The reads were counted using HTSeq v. 2.0.2 (Putri *et al*., 2022). For further analysis the samples were combined in a single table.

Analysis of differential gene expression was performed with DESeq2 v.3.16 (Love *et al.,* 2014) in R and R Studio v.2022.07.02+576 (R Core Team, 2018; R Studio Team, 2020). The data was filtered to contain only genes with at least ten reads in at least four samples. For a preliminary quality assessment, the data was variance stabilised using the function varianceStabilizingTransformation from the DESEq2 package v.1.12.3 (Love *et al.,* 2014). Then, a Principal Component Analysis (PCA) was plotted with the function plotPCA from the package DESeq2. Additionally, a heatmap was plotted with the variance stabilised counts data using pheatmap v.1.0.12 (Kolde, 2010) for visualisation of the sample quality. Differential gene expression analysis was performed with default settings of the function DESeq2, with an adjusted p-value of p<0.05. Plots were produced in R (R Core Team, 2018) with the packages ggplot2 v.3.3.6 (Wickham, 2016) and reshape2 v.1.4.4 (Wickham, 2007). A BlastX search against the nrProt database (v. January 2022) was conducted which resulted in annotations for 86% of genes. The raw data of all samples sequenced can be found at ENA (study number: PRJEB55616).

### 2.3 Gene enrichment analysis

The Interproscan database (v. January 2022) was used to obtain Gene Ontology (GO) information for all proteins of the *C. riparius* genome. A functional enrichment analysis in the category “biological function” was performed with all significant DEGs, and with all up- and downregulated DEGs separately. The analysis was conducted with the package topGO v.2.24.0 (Alexa & Rahnenfuhrer, 2022) using the ‘weight01’ algorithm and a p-value cut-off of <0.05 (Fisher’s test). For reasons of statistical relevance, we only considered GO terms based on at least five annotated genes.

### 2.4 Fitness test

For the remaining ten replicates per treatment, the number and sex of emerged individuals were recorded daily to calculate mortality, sex ratio and median time to emergence of 50% of females of the population (EmT50). Emerged individuals were transferred from bowls to cages for mating and oviposition. Egg ropes were collected until the end of the experiment. Fertility was measured according to Foucault et al. (2019), consisting of the number of egg ropes where at least 50% of the larvae hatched, divided by the number of females. Finally, mortality, female EmT50 and fertility values were integrated in a simplified Euler-Lotka calculation to assess PGR. This last measure is a composite fitness measure where each parameter is not only summarized but weighted according to its impact on the population (Nemec et al., 2013). Variances of the treatment groups were compared with F-tests on equal variances using the function var.test from the package stats v.3.6.2 in R. A Bayesian implementation of a t-test to was used to check for differences in fitness components and resulting PGR between control and ALAN treatment (R library BayesianFirstAid, Bååth, 2014).

To assess the potential long-term demographic consequences of the observed differences in daily population growth rates between control and ALAN treatment, we calculated the ratios of the cumulative population sizes of two populations growing at the respective rates for 200 days as a geometric progression.

It was noticeable that the variance of some life cycle parameters appeared to be much lower under ALAN than under control conditions for developmental time, fertility and PGR (Figure 5B-D). This impression was confirmed by F-tests on equal variances that confirmed (marginally) significant differences for all parameters

### 2.5 In vivo reactive oxygen species (ROS) level test

As the transcriptomic analysis suggested that oxidative stress might increase in the larvae in response to ALAN, we tested this hypothesis in a *post-hoc* experiment by measuring ROS levels *in vivo*. ALAN and control conditions were set to the same settings as in the life-cycle test. Per treatment group, 20 L3-stage larvae were collected from the lab culture and placed in 24-well plates. The measurements were carried out as described in Rigano et al., 2024. Well plates were filled with 2.5 mL medium as described in Foucault et al., 2019.

ROS products were identified with CellROX Orange (Thermo Fisher cat. no. C10443). CellROX is a cell-permeable oxidative stress reagent and may be used for *in vivo* ROS measurements. It is non-fluorescent in its reduced state. Upon oxidation by ROS, the reagent emits a bright orange fluorogenic signal at 545/565 nm. The signal from CellROX orange is localised within the cytoplasm and can detect 5 different ROS types (hydrogen peroxide, hydroxyl radical, nitric oxide, peroxynitrite anion, and superoxide anion).

After placing larvae on the well plates, 0.75 μL of CellROX Orange was used per larva. Well-plates were placed in a climate chamber with ALAN and control conditions as described in chapter 2.1. Larvae were subjected to 24 h of treatment, then the well plates were placed in a styrofoam box to avoid any temperature change effects. ROS levels were measured in live larvae with ZEISS Axio Imager 2 under 10 x magnification. Images were taken with AxioVision Rel. v.4.8. For fluorescence imaging, an HXP 120C fluorescence lamp was used at maximum intensity (Item Number: 423013-9010-000). Fluorescence images of the larvae were captured using the “43 HE” filter set (BP 550/25 HE, FT 570 HE, BP 605/70 HE, Item Number 489043-9901-000) with a 1 s exposure time.

This filter set excites blue light at approximately 550 nm, transmits emitted red fluorescence above 570 nm, and filters out residual blue excitation light, allowing only red fluorescence around 605 nm to be detected. Fluorescence field images were analysed using ImageJ Fiji v. 2.15.0. Images were imported into ImageJ as a sequence and converted to 8-bit grayscale from RGB colour to eliminate colour variations and focus solely on light intensity. A consistent threshold of 23 was applied to all images. The measure function was used to calculate the mean intensity value for each image. A Bayesian t-test was employed to identify significant differences in relative fluorescence levels between the control and ALAN treatment groups using the R library BayesianFirstAid (Bååth, 2014).

### 2.6 Observation of development

From the transcriptomic analysis we predicted that the onset of metamorphosis was earlier under ALAN, though the life cycle test showed higher EmT50 in this group, i.e., midges emerged later. We hypothesised that ALAN increased the time spent in metamorphosis. To test our hypothesis, we performed a second posthoc experiment to clarify the changes in onset and duration of metamorphosis due to ALAN. The experimental conditions were identical to the life cycle test. We collected 40 freshly hatched larvae from five pooled egg clutches from the laboratory culture and divided them among two climate chambers which each contained one replicate bowl with 20 individuals. The experimental conditions were identical to the life cycle test and one group was subjected to ALAN conditions and one to control conditions. Once the larvae had reached the L4 stage we transferred them to a new bowl which contained only a handful of sediment so that we could better observe their development. We installed a wildlife camera (Coolife H881 PLUS) above the bowls which took one picture every 30 minutes. Once all midges had emerged, we analysed the images. Unfortunaly the image quality was disparate and the analysis only possible for the determination of the onset of ecdysis. We performed a Bayesian implantation of a t-test (R library BayesianFirstAid, Bååth, 2014) to check for differences in the onset of moulting between the ALAN and control group.

## 3. Results

### 3.1 Gene expression analysis

Sequencing resulted in 188,760,770 raw paired reads (28.3Gb) for the control, and 159,468,341 paired reads (23.9Gb) for the ALAN treatment. Filtering for genes with at least 10 mapped reads left 11,715 of the 13,449 annotated genes in the genome assembly. In total, 1,564 (11.6% of all annotated genes) significantly Differentially Expressed Genes (DEGs) were identified between the two treatments. Of these, 786 genes were upregulated and 778 genes were downregulated under ALAN. The annotated DEG with the highest expression increase was calphotin-like, which is linked to photoreception (Ballinger et al., 1993). Amongst other upregulated DEGs, we found a nuclear hormone receptor (NHR), a nuclear factor (NF) and the enzyme E3 ubiquitin-protein ligase HUWE1, all of which were linked to the regulation of the circadian clock. Six out of eight methyltransferases were downregulated and two histone acetyltransferases (HATs) were upregulated. Both transferases play a role in epigenetic regulation.

DEGs involved in development included three genes of the juvenile hormone pathway, four genes linked to the moulting hormone ecdysone, six dead box proteins, two upregulated pupal cuticle proteins, chitinase 1, chitin deacetylase 1, chitin synthase chs-2 isoform X3 and ten cytochrome P450 isoforms. The DEG circadian clock protein daywake-like was downregulated. The NCBI database reports juvenile hormone hemolymph binding protein (hJHBP) as a protein ortholog for this gene in *Drosophila* (https://www.ncbi.nlm.nih.gov/gene/101743480). All DEGs of the juvenile hormone pathway were downregulated, while genes related to ecdysone were upregulated, with the exception of ecdysone-20-monooxygenase (figure 2d), which was downregulated. We found eight ATPases which are linked to catabolism.

**Figure 1:**
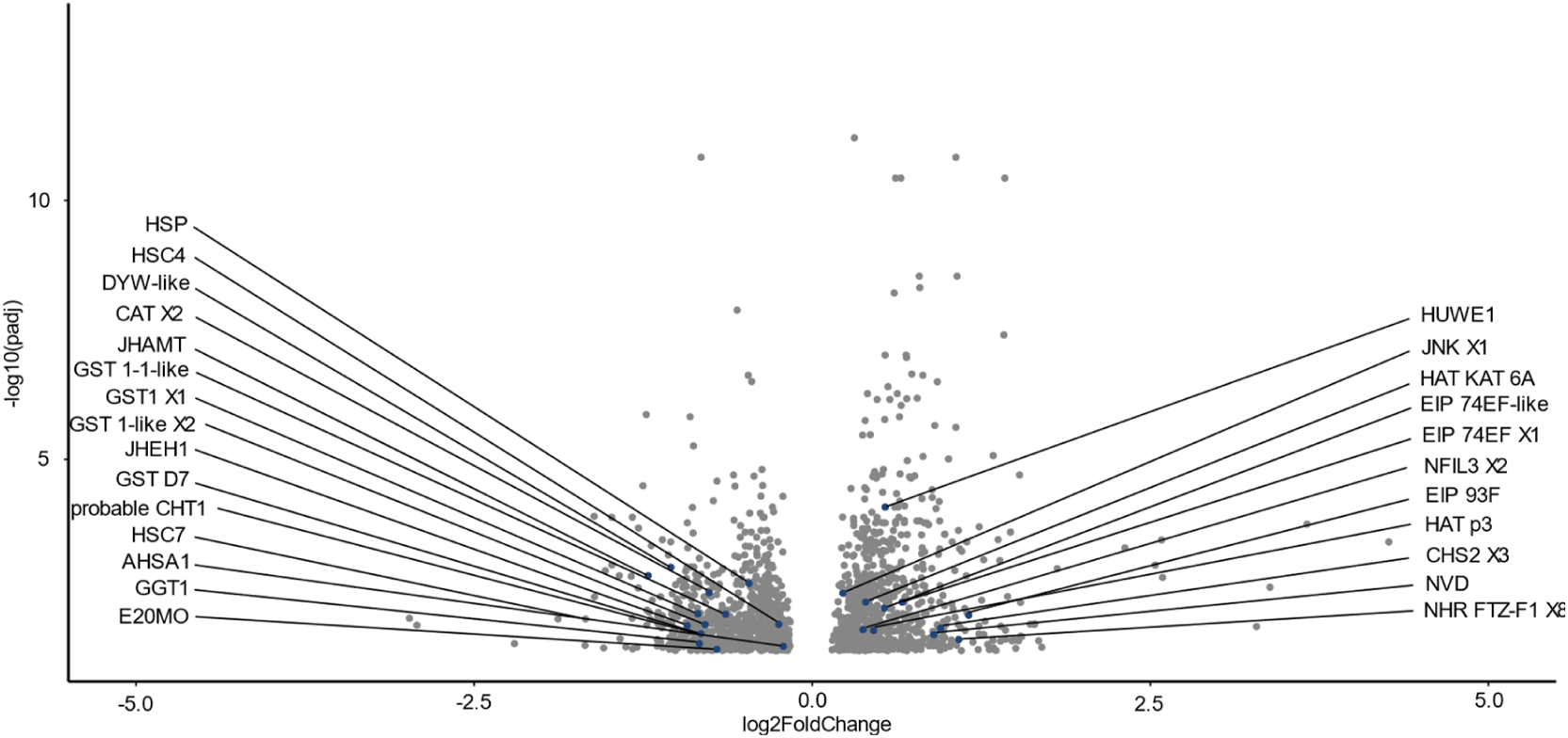
Volcano plot of all significantly differentially expressed genes (DEGs) with padj<0.05. Labelled DEGs are candidate genes which were investigated further. HUWE1: E3 ubiquitin-protein ligase HUWE1, dyw: circadian clock-controlled protein daywake-like, JHAMT: juvenile hormone acid O-methyltransferase, HSP: heat shock protein 27-like, CAT X2: catalase isoform X2, JNK X1: stress-activated protein kinase JNK isoform X1, HAT KAT6A: histone acetyltransferase KAT6A, EIP 74EF X1: ecdysone-induced protein 74EF isoform X1, EIP 74EF- like: ecdysone-induced protein 74EF-like, EIP 93F: Putative uncharacterized protein DDB_G0277255 isoform X1 (KEGG Orthology code K20015: ecdysone-induced protein 93F), GST1 X1: glutathione S-transferase 1 isoform X1, GST 1-1-like: glutathione S-transferase 1-1- like, HSC4: heat shock 70 kDa protein cognate 4, GST 1-like X2: microsomal glutathione S-transferase 1-like isoform X2, JHEH1: juvenile hormone epoxide hydrolase 1, CHS2 X3: chitin synthase chs-2 isoform X3, HAT p300: histone acetyltransferase p300, NFIL3 X2: nuclear factor interleukin-3-regulated protein isoform X2, NVD: cholesterol 7-desaturase nvd, GST D7: glutathione S-transferase D7, probable CHT1: probable chitinase 10, NHR FTZ-F1 X8: nuclear hormone receptor FTZ-F1 isoform X8, GGT1: glutathione hydrolase 1 proenzyme-like, AHSA1: activator of 90 kDa heat shock protein ATPase homolog 1, E20MO: ecdysone 20- monooxygenase.

**Figure 2:**
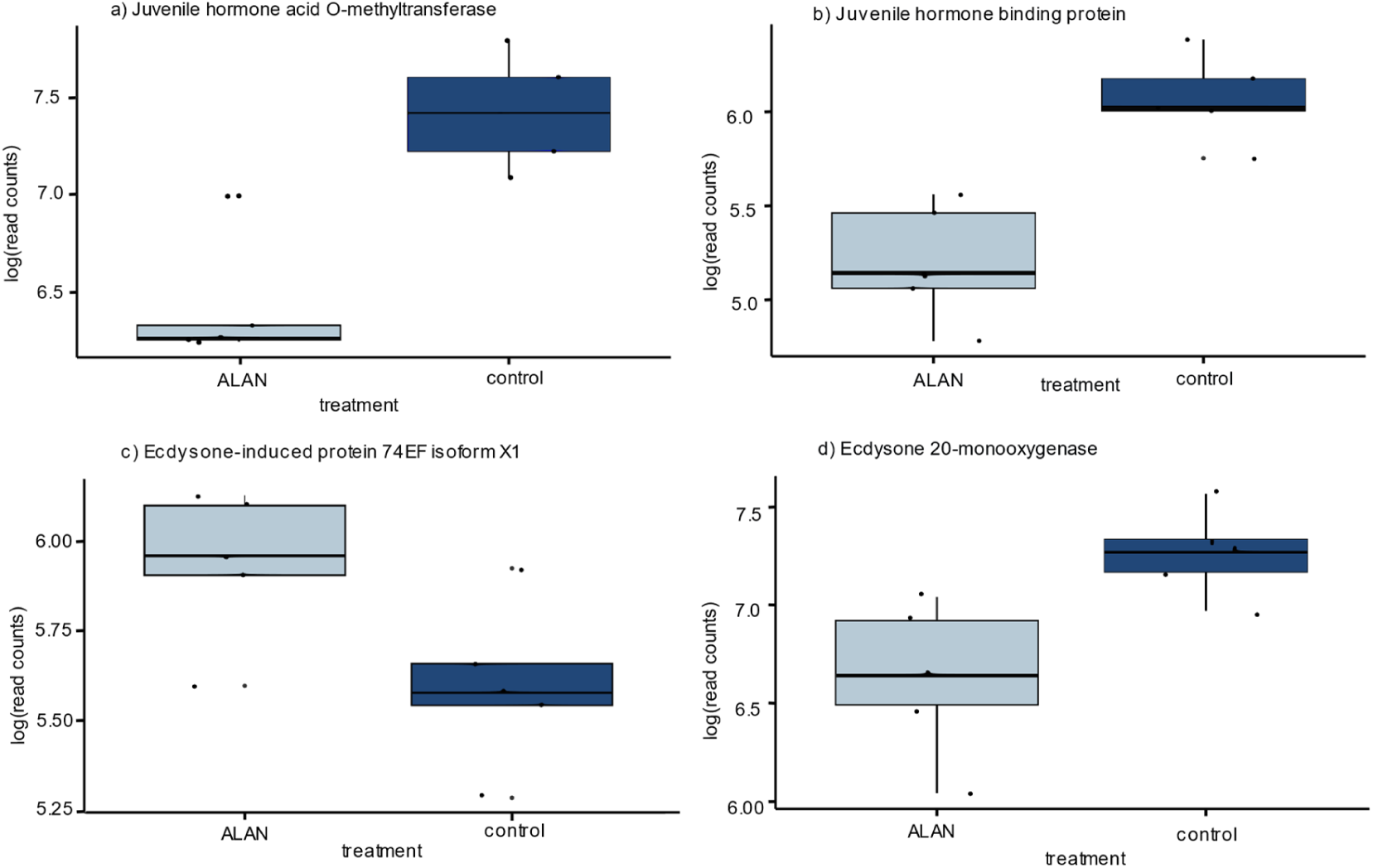
Expression patterns of the significantly DEGs linked to juvenile hormone and ecdysone which control molting and show that ALAN altered developmental processes in the chironomids. a) Juvenile hormone acid O-methyltransferase (base mean=1152.71; Log2 Fold Change (L2FC)=-1.21; padj<0.01**) was downregulated under ALAN. b) Circadian clock-controlled protein daywake-like was downregulated in the ALAN treatment (base mean=305.65; L2FC=-1.046, padj<0.01**). This gene encodes the protein juvenile hormone hemolymph binding protein in *Drosophila melanogaster* (https://www.ncbi.nlm.nih.gov/gene/101743480). c) Ecdysone-induced protein 74EF isoform X1 was upregulated under ALAN (base mean=331.15; L2FC=0.67; padj<0.01**). d) Ecdysone 20-monooxygenase was downregulated in the ALAN treatment (base mean=1098; L2FC=-0.69; padj<0.05*). P-value thresholds were defined as: p<0.05*=0.05>p>0.01; p<0.01**=0.01>p>0.001; p<0.001***).

We identified twelve downregulated DEGs that are related to the antioxidant defence system, namely a catalase isoform, six glutathione S-transferases, three heat shock proteins and an activator of 9 kDa heat shock protein ATPase homolog 1. A peroxidase-like isoform, also belonging in this functional context, was upregulated. A list of all candidate genes can be found in the Supplementary Material S1.

### 3.2 GO term enrichment analysis

In total 4 significantly enriched functions were identified in either up- or downregulated genes (Table 1). Upregulated GO terms were, in order of descending number of significant genes, Rho protein signal transduction, ion transport and potassium ion transport. The downregulated GO term was glutathione metabolic process.

**Table 1:** Significantly enriched biological functions based on genes differentially expressed between the ALAN treatment and control. Separate GO analyses were performed for up- and downregulated genes. Annotated: number of genes with according GO annotation in the reference set. Significant: number of genes with according GO annoation in the test set. p-value: result of Fisher’s exact test, based on gene counts.

| GO ID | GO Term | Annotated | significant | Expected | p-value |
| --- | --- | --- | --- | --- | --- |
| *upregulated* |  |  |  |  |  |
| GO:0007266 | Rho protein signal transduction | 116 | 9 | 3.79 | 0.02 |
| GO:0006811 | ion transport | 28 | 4 | 0.92 | 0.011 |
| GO:0006813 | potassium ion transport | 5 | 2 | 0.27 | 0.026 |
| *downregulated* |  |  |  |  |  |
| GO:0006749 | glutathione metabolic process | 5 | 2 | 0.11 | 0.0041 |

### 3.3 Effect of ALAN on ROS level

After one night under ALAN conditions, the Bayesian analysis indicated a significant rise in oxidative stress compared to the control group. A difference of means of 2 relative fluorescence units under ALAN compared to control conditions was found (95% highest density interval (HDI): −0.82 to 4.7) with a posterior probability of 91.7% that oxidative stress was higher under ALAN. The effect size was found to be of small to medium magnitude (Cohen’s d = 0.46. (Figure 3).

**Figure 3:**
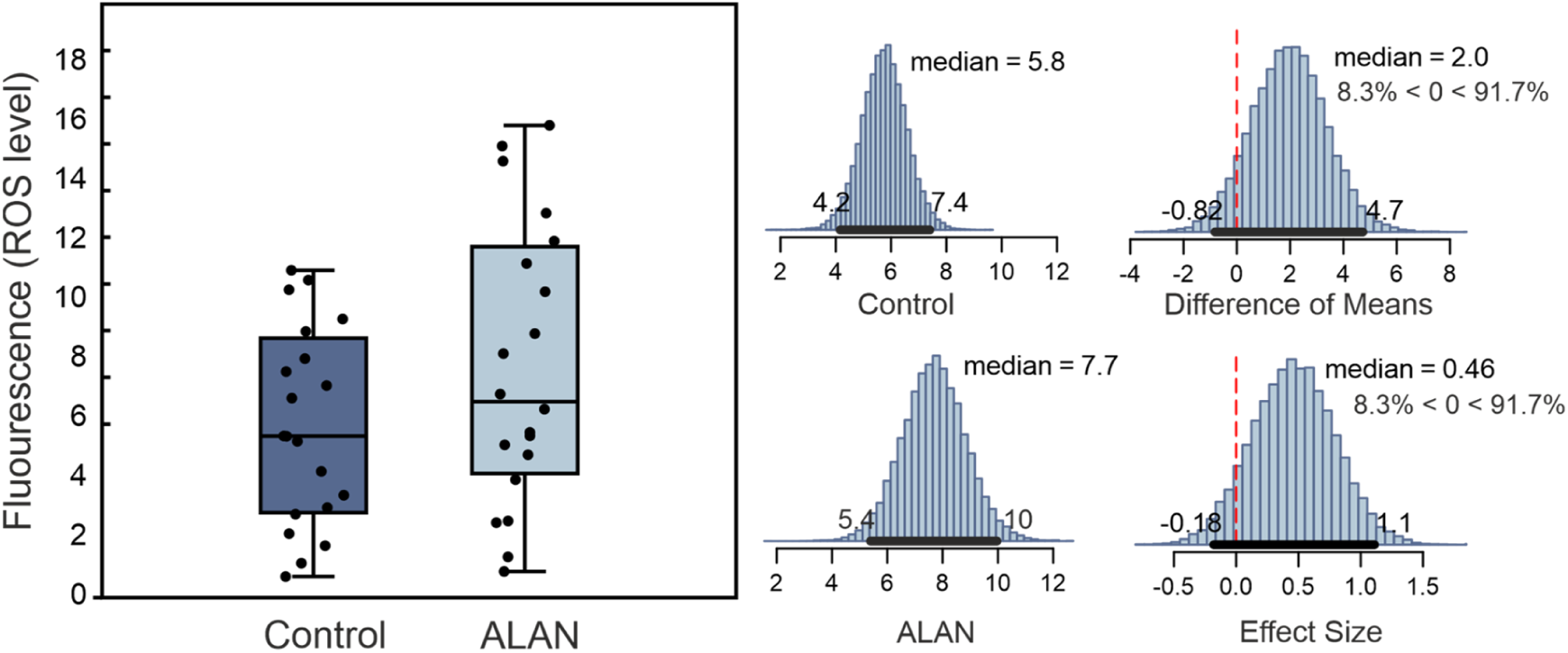
Comparison of ROS level measured as relative fluorescence intensity after 24 hours under ALAN and control conditions. The histograms in the middle show the Bayesian posterior estimates for the medians in the control (top) and ALAN (bottom) group. The difference of means (top right) has a 91.7 % posterior probability of being larger in the ALAN group. The median effect size has the same posterior probability of being larger than zero (bottom right).

### 3.4 Observation of development

The camera images revealed strong evidence for earlier ecdysis caused by ALAN. The Bayesian analysis showed a difference of mean of −2.2 days (95% HDI: −4.3 to −0.18) with a posterior probability of 98.1% that ecdysis commenced earlier under ALAN. This was supported by a very large effect size (Cohen’s d = −1.5) (Figure 4).

**Figure 4:**
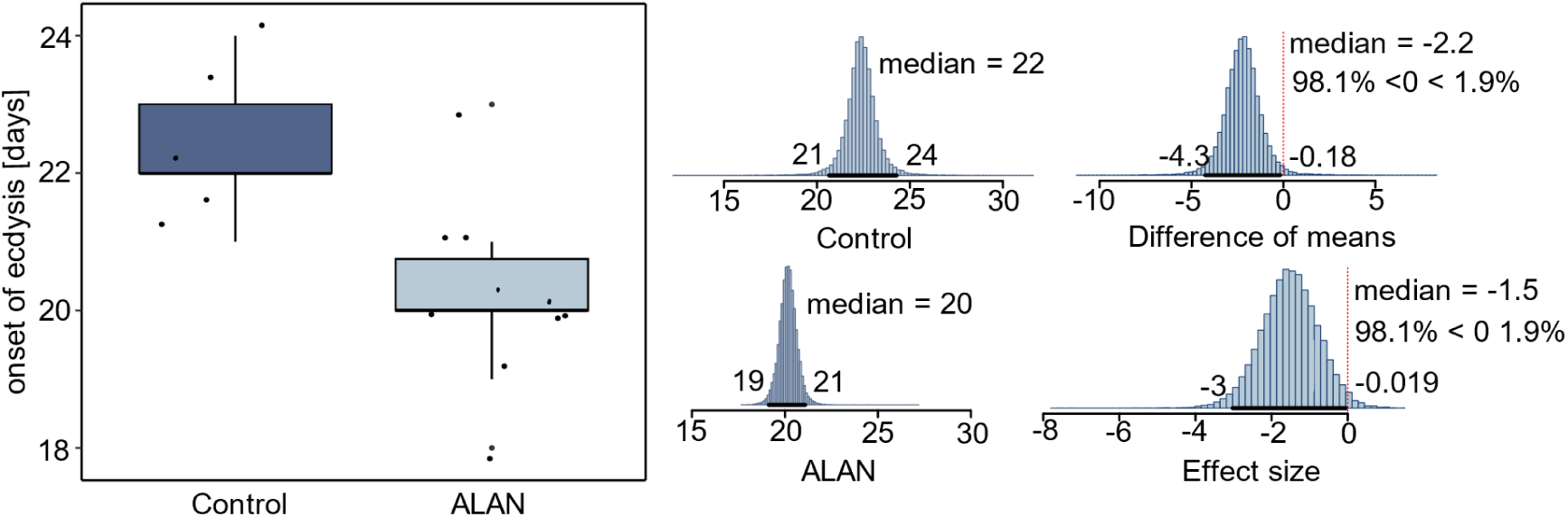
Time difference until onset of pupation under ALAN and control conditions. There was strong evidence (98.1%) for earlier ecdysis (2.2 days) under ALAN (95% HDI = −4.3 – 0.18) with very large effect size (Cohen’s d = −1.5). Difference in days to pupation between ALAN and control groups. The histograms in the middle show the Bayesian posterior estimates for the medians in the control (top) and ALAN (bottom) group. The difference of means (top right) has a 98.1 % posterior probability of being smaller in the ALAN group. The median effect size has the same posterior probability of being smaller than zero (bottom right).

### 3.5 Fitness test

Out of 600 first instar larvae, 564 adults emerged in total (94%). The differences in mortality between Control and ALAN treatment were very low, with no evidence for any relevance (Figure 5A).

There was evidence for a prolongation of 0.27 (95% HDI: −1.2 to – 0.67) days in female EmT50 with a posterior probability of 72.8% and small effect size (Cohen’s d = −0.28) under ALAN conditions (Figure 5B).

Very strong evidence and large effect size (Cohen’s d = 1.2) were found with a posterior probability of 98.8% that ALAN caused lower fertility. On average, an individual produced 34 (95% HDI: 5.2 to 61) fewer viable eggs under ALAN conditions than in the control group (Figure 5C). Integrating the fitness components into a daily population growth rate showed decisive evidence (posterior probability: 100%) for a strong reduction (−0.031/day, 95% HDI: 0.018 to 0.044) of immense effect size (Cohen’s d = 2.4, Figure 5D) under ALAN conditions.

It was noticeable that the variance of some life cycle parameters appeared to be much lower under ALAN than under control conditions for developmental time, fertility and PGR (Figure 5B-D). This impression was confirmed by F-tests on equal variances that confirmed (marginally) significant differences for all parameters (F = 2.84 – 5.94, p = 0.051 - 0.014).

**Figure 5:**
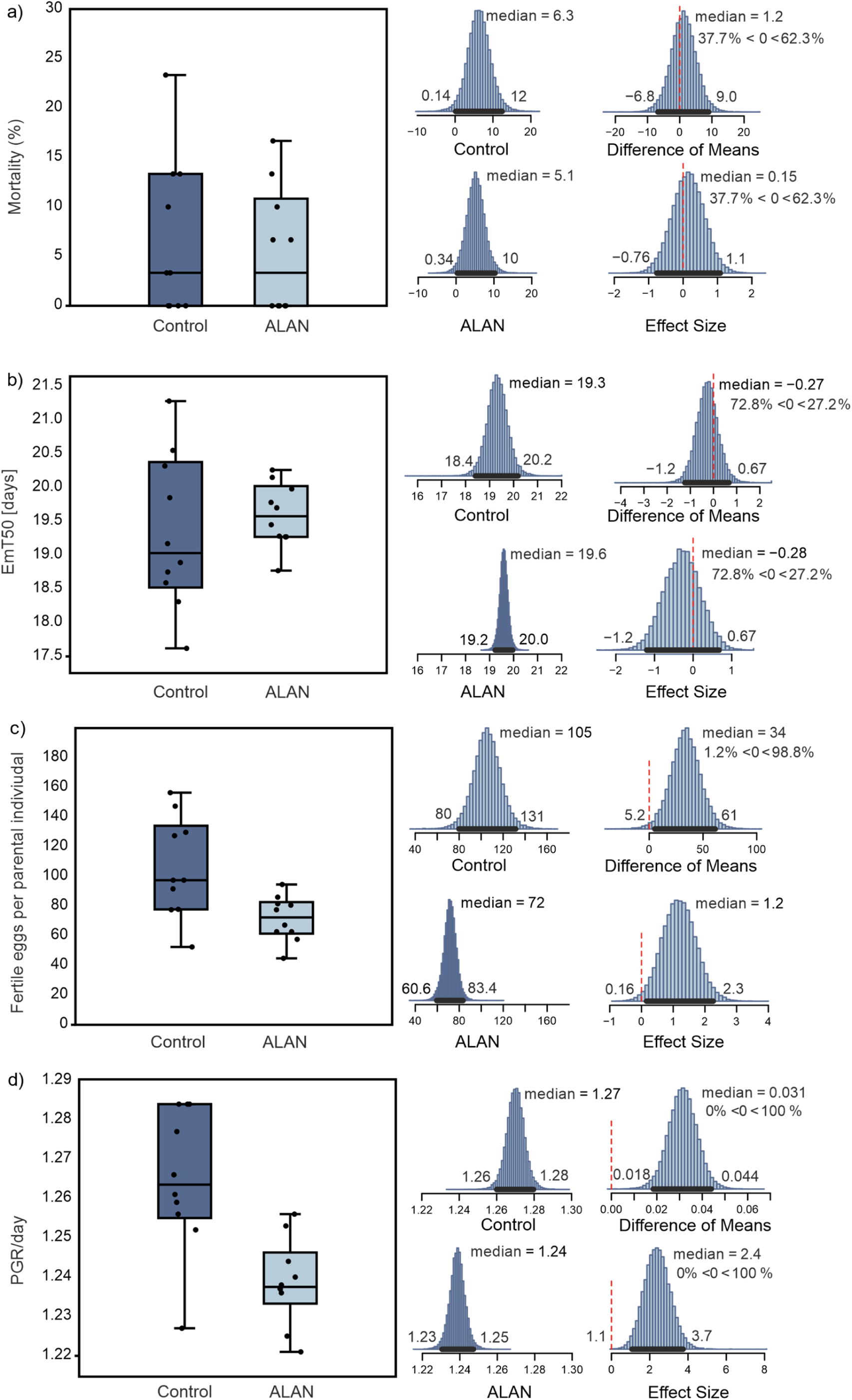
There was no evidence for any relevant difference in mortality between control and ALAN treatment (95% HDI: −6.8 to 9.0). There was some evidence for prolonged EmT50 of 0.28 (95% HDI: −1.2 to 0.67) days with small effect size (Cohen’s d = −0.28) as indicated by a posterior probability of 72.8%. Fertility was reduced by 34 (95% HDI: 5.2 to 6.1) fewer viable eggs under ALAN conditions. Integrating the fitness components into a daily population growth rate showed decisive evidence (100%) for a strong reduction (−0.031/day, 95% HDI: 0.018 to 0.044) of immense effect size (Cohen’s d = 2.4) under ALAN conditions.

The difference in daily growth rate was between 1.8 and 4.4 percent which resulted quickly in substantial differences in potential population size (Figure 6). Under ALAN conditions, a population would grow to half the size of a respective control population after 28 days (95% HDI: 18 to 67 days). After 200 days, the approximate length of the annual reproductive period in the field, a population under ALAN conditions would have produced only about 1% of the individuals compared to a population without artificial light exposure at night.

**Figure 6:**
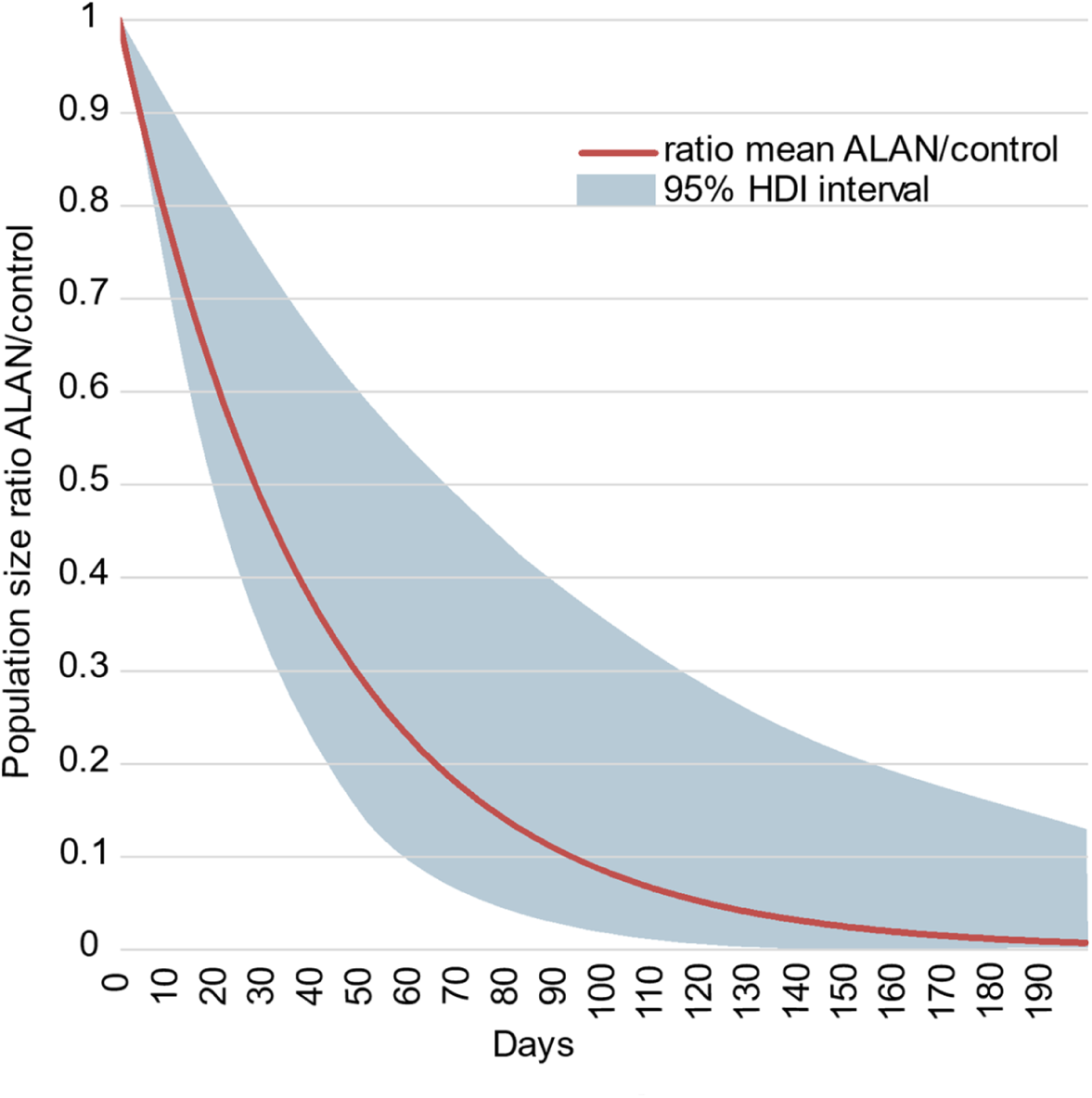
Trajectory of potential population size ratios under ALAN (PGR 1.24/d, 95% HDI: 1.23 to 1.25) and control (PGR 1.27/d, 95% HDI: 1.26 to 1.28) daily population growth rates for 200 days (i.e., the approx. annual reproduction period). This PGR difference of 0.031 (95% HDI: 0.018 to 0.044) per day implies that the ALAN population attains half the size of the control population after 28 days and reaches only 1% of an undisturbed population after 200 days.

## 4. Discussion

### ALAN substantially changed gene transcription patterns

The exposure of *C. riparius* to light at night altered the gene expression patterns on a broad scale. A minimum of 11.6% of all annotated genes in the genome were substantially either up- or downregulated due to the nightly light exposure of 100 Lux, which suggested a comprehensive systemic response to this environmental change. This interpretation is supported by a substantial number of differentially regulated genes with functions in circadian rhythm, development, and stress response.

### Expression changes of circadian rhythm and developmental genes predicted an earlier onset of metamorphosis

The rhythmic oscillations of circadian clock gene expression are maintained and synchronised by the natural daily cycling of light and dark periods (Abe *et al.,* 2022, Carmakian & Boivin, 2009). These genes in turn control the timing and speed of developmental processes, such as moulting and eclosion (Franco *et al.,* 2017, in Brady *et al.,* 2021, Feng & Lazar, 2012, Tomioka & Matsumoto, 2010, Saunders, 2002). We found several genes related to circadian rhythmicity among the DEGs. Differential expression of a nuclear hormone receptor (NHR) and a nuclear factor (NF), as well as two histone acetyltransferases as found here were linked in previous studies to the circadian rhythm, as well as developmental processes in *Drosophila* and vertebrates (Qiu *et al.,* 2023, Burris, 2015). An experiment with ants showed that inhibition of histone acetyltransferases caused a loss of circadian rhythmicity, disabling their ability to adjust their activity to rhythms of light and dark periods (Libbrecht *et al.,* 2020). Although we did not explicitly measure activity patterns in our experiment, it is possible that the differential expression of these genes had a similar effect.

Next to indications for altered circadian rhythmicity we also found that ROS levels were increased in the ALAN group. There is evidence for a bidirectional link between circadian rhythmicity and oxidative stress in zebrafish (*Danio rerio*). Genes related to antioxidant and immune responses show circadian rhythmicity and therefore, sensitivity to stressors differ depending on the time of day (Zheng *et al.,* 2017). On the other hand, environmental stressors may disrupt the endogenous circadian clock (Doria *et al.,* 2018). Disturbance of the circadian rhythm caused by the ALAN treatment could have therefore increased the midges’ sensitivity to ROS and thus, increasing oxidative stress damage.

DEGs linked to developmental processes included genes related to the juvenile hormone (JH) and ecdysone. JH and ecdysone determine the onset of moulting and metamorphosis (Qiu *et al.,* 2023; Jindra *et al.,* 2015, *Bond et al*., 2010). JH prevents metamorphosis until the larva has reached an appropriate size (Qiu *et al.,* 2023; Daimon *et al.,* 2015). JH acid methyltransferase (JHAMT) is part of the JH biosynthesis pathway (Shinoda & Itoyama, 2003) and was downregulated in this experiment (Figure 2a). The protein ortholog of the DEG, circadian clock protein daywake-like is the juvenile hormone hemolymph binding protein (hJHBP), https://www.ncbi.nlm.nih.gov/gene/101743480). hJHBP transports JH from the *corpora allata*, where it is produced, to the target tissues and shields it from carboxylesterases which would convert the JH molecule to its inactive form (Zalewska *et al.,* 2009; Kramer *et al.,* 1976). Hence, a down-regulation of hJHBP, as it was observed under ALAN, could cause further degradation of active JH. This indicated that under ALAN, the transition from larvae to pupation was already ongoing. This interpretation was supported by the expression patterns which concur with those of the oriental fruit fly *Bactrocera dorsalis* during pupariation (Chen et al., 2018): We observed an upregulation of the pupal cuticle proteins 2-like and 27-like, the downregulation of chitinase 1 and cytochrome P450. These proteins play important roles during metamorphosis and are activated during the transition to the adult insect (Shahin et al., 2018).

The NHRs were previously discussed with regard to their role in the circadian clock. The NHR FTZ-F1, which was upregulated under ALAN, also regulates pupation in the moth *Helicoverpa armigera* (Zhang *et al.,* 2021). In their experiment Zhang *et al*. (2021) could show that knockdown of FTZ-F1 repressed the expression of metamorphosis-inducing ecdysteroidogenesis genes and induced the transcription of JH biosysnthesis genes in *H. armigera*. We found reverse expression patterns with upregulation of FTZ-F1 and downregulation of genes related to the JH biosynthesis. Two other studies found similar effects in silkworms when the circadian clock was manipulated (Daimon *et al*., 2015, Qiu *et al.,* 2023), leading to either premature or delayed metamorphosis. The authors concluded that expression of JH is affected by a circadian clock transcription factor through regulation of JHAMT-like (Qiu *et al*., 2023). These findings further support a link between the circadian clock and the regulation of developmental hormones in chironomids.

The above discussed evidence from the gene transcription analysis for an earlier onset of metamorphosis was at first sight contradictory to our findings in the life cycle experiments, which indicated an increase in developmental time caused by the ALAN treatment. Since we confirmed that pupation begins earlier under ALAN in an additional experiment, we could show that metamorphosis is in fact longer in this group and thus refute the seeming contradiction.

### ALAN synchronised and streamlined developmental processes

The DEGs encoding ATPases indicated changes in energy metabolism under ALAN. During the maturation of the pupae, energy metabolism is increased for the differentiation of organs (Ozerova & Gelfand, 2022), thus providing further evidence for differential but streamlined development in the two treatment groups. In addition, the transcriptomic analysis showed genes linked with moulting in the ALAN group, indicating an earlier onset of pupation in this group.

To test our hypothesis, we captured the developmental progress of midges in ALAN and control chambers with the use of wildlife cameras. We were able to confirm an earlier onset of pupation in the ALAN group. While midges under ALAN emerge later as seen in the fitness test, metamorphosis begins earlier in this group as proposed by our gene expression analysis. Therefore, the duration of metamorphosis is increased due to the effect of ALAN. A previous study (Doria *et al*., 2022) observed longer developmental time of *C. riparius* with longer day length and suggested that adaptation to seasonal photoperiodic variation might be the cause of this. Constant light, viewed as an extreme photoperiodic condition, could therefore cause a beneficial adaptive response of prolonged development.

### Oxidative stress-related genes

Fertility was the parameter that most influenced PGR results, being the observed fitness loss mainly due to a reduced number of egg ropes per female, indicating a stress induced fertility impairment (Sales *et al.,* 2021), more visible at constant light conditions. Among the DEAD-box proteins (DDX) which were downregulated under ALAN, knockdown of DDX47 was linked to seized ovarian development and delayed moulting in the planthopper *Laodelphax striatellus* (Ma *et al*., 2024).

Although we found no research on the effect of light duration on fertility, temperature stress has been shown to impair the development of reproduction in the flour beetle *Tribolium castaneum* where the effect of stress differed between developmental stages with the highest impact during the pupal stage (Sales *et al*., 2021).

Moreover, photoperiod acts as a reliable seasonal cue. Longer day lengths indicate a later time in the year, hence a shorter development through increased growth rate and reduced size at maturity could be shown for C. riparius, butterflies and caddis flies (Doria *et al*., 2021, Dmitriew, 2011). The prolonged photoperiod may therefore either trigger a stress response, or it may cause an adaptive response to the season.

Reactive oxygen species (ROS) are molecules which are formed as by-products during cellular metabolism and that have various functions involving their ability to oxidate other molecules (Jakubczyk *et al*., 2020). Several mechanisms exist for cells to maintain redox homeostasis, however, excessive levels of ROS tip the levels in favour of oxidation, causing oxidative stress which leads to damage of cells (Jakubczyk *et al*., 2020).

A glutathione S-transferase (GST) and three heat shock proteins (HSPs) were downregulated in the treatment group, whereas a peroxidase was upregulated. GST is involved in the removal of reactive oxygen species (ROS) from cells (Yuan *et al*., 2013), so a downregulation of these two enzymes causes a build-up of ROS. Moreover, fluctuating levels of CAT and GT are also observed during the development of various organisms. Previous studies showed a down-regulation of antioxidant enzymes during development of amphibians and *Drosophila*, where the resulting peak in hydrogen peroxide after a drop in GT, among other antioxidants, was needed for apoptosis in the remodelling of the organism during metamorphosis (Menon & Rozman 2007, Macho *et al.,* 1997; Bewley *et al.,* 1983). In the moth *Achaea Janata*, CAT peaked during the 5^th^ instar stage and dropped during pre-pupal and pupal stages (Pavani *et al*., 2015). Therefore, it is not clear from the transcriptome analysis whether the observed differential expression of antioxidant enzymes is linked to the developmental stage of the organisms, i.e., onset of metamorphosis, a stress response, or even both.

A GO term search of upregulated DEGs resulted in potassium ion transport, among others. There is evidence that potassium channels protect cells from mitochondrial oxidative stress which indicated an increase of oxidative stress under ALAN (Szweczyk *et al*., 2009, Slocinska *et al*., 2013). A GO term search of downregulated DEGs resulted in glutathione metabolic process. Overall, even though an increase of ROS can occur naturally during the development of holometabolous insects, the observed downregulation of the antioxidant defence system under ALAN should lead to an (additional) increase of ROS.

To test this hypothesis, we performed an additional 24 hour experiment in which we subjected individuals of the same age to ALAN and control conditions for 24 hours and then measured ROS, thereby removing the influence of potential differences in developmental stage. Assuming that the measured ROS concentration under control conditions represents the base level in L4 chironomid larvae, the increase in intracellular ROS level should therefore be caused by the ALAN treatment, potentially caused by the observed downregulation of the enzymatic oxidative stress response relative to the control conditions.

### Fitness test reveals dramatic cumulative population decline

The transcriptomic analysis revealed differential expression of genes related to the circadian rhythm, development and stress. The life cycle experiment confirmed measurable effects on development and fitness impairment. Individuals under ALAN emerged later and within a smaller time window than in the control group. Moreover, the onset of emergence between sexes was temporally spaced further apart than in the control group.

A strong fitness reduction was observed under ALAN as fertility and population growth rate (PGR) were significantly reduced, although there was no difference in mortality between groups. Plotting the trajectory of population size given the calculated PGR under ALAN shows a dramatic decrease over a vegetation period of 200 days. After only 27 days, the population is already reduced to half of its original size. While there is evidence that ROS directly reduces fertility through sperm damage (Reinhardt & Ribou, 2013), fecundity may also be reduced through lower body mass caused by higher activity levels. In previous studies a positive relationship between female body mass and fertility could be observed for *Chironomus riparius* (Péry *et al*., 2002, Taylor et al. 1998) and *tentans* (Sibley *et al*., 2001), while a negative effect of ALAN on body mass has been shown in the moth *Apamea sordens* (Grenis *et al*., 2023). Moreover, in numerous Lepidopterans, the number of eggs laid as adults is positively correlated with body mass in female larvae (Loewy *et al.,* 2013). As previously discussed, HATs which were found in our analysis have been shown to interfere with ants’ activity patterns (Libbrecht *et al.,* 2020), and could have led to prolonged activity of chironomids under ALAN, thus reducing body mass in females. In addition, a previous study found that fertility increased with longer dark periods (Doria *et al*., 2021). This points at two possible explanations: i) constant lighting causes females to be more active, leaving fewer resources to the development of fecundity, or ii) development of fertility could be a process limited to dark periods.

### Detrimental fitness effects of ALAN could influence demography and ecological interactions

At first sight, the effects of ALAN on the population growth rate at hatching with a decrease by about 3% relative to the control seemed to be limited, as populations under both treatments would still grow more than 120% per day. However, when projected over 200 days, the approximate period of *C. riparius* reproduction in temperate areas (Waldvogel *et al*., 2018), the cumulative effect of this difference on population size would be dramatic. Regardless of the extent or cause of expected post-hatch mortality, a population growing at the ALAN rate would cumulatively over a growing season reach only 1% of the size of a population without this anthropogenic influence. Given the central role in freshwater and terrestrial food webs (Schulz *et al*., 2024) and the huge population sizes of Chironomids (Pinder, 1986), ALAN could locally alter community compositions of insects and impact the function of entire ecosystems through cascading effects (Hirt *et al*., 2023). Moreover, if the results found here can be extended to other insects, ALAN could be a yet underestimated driver under the many negative anthropogenic impacts on insect demography, leading synergistically to the observed insect decline (Wagner *et al*., 2021).

### Transcriptomic analysis accurately predicts physiological changes

Next to the ecological effects of ALAN on populations, we were able to show that transcriptomic analyses are able to accurately predict physiological changes. From the significantly DEGs we predicted that 1) ALAN causes increased levels of ROS in the midges, and that 2) the onset of pupation is earlier and synchronised under ALAN. In the two posthoc experiments we could confirm these predictions and therefore conclude the usefulness of transcriptomic analyses in predicting effects before they can even be observed in experiments.

## 5. Conclusion

In this study we showed that artificial light at night (ALAN) causes oxidative stress in *C. riparius* and alters development through earlier onset of metamorphosis and prolonged duration thereof. We observed a severe and swift reduction in population size with a reduction to 1% after 200 days of the predicted population. Considering the large and increasing occurrence of ALAN, it could constitute a major factor in the observed insect decline. This study adds to the increasingly relevant topic of light pollution and demonstrates the viability of transcriptomic analyses to predict higher-level phenotypic changes.

## CRediT authorship contribution statement

**Linda Eberhardt**: Writing – original draft, Visualization, Validation, Methodology, Investigation, Formal analysis, Data curation. **Halina Binde Doria**: Conceptualisation, Investigation, Formal analysis, Supervision, Writing – review & editing, Data curation. **Burak Bulut**: investigation, formal analysis, writing – review and editing. **Barbara Feldmeyer**: formal analysis, writing – review and editing **Markus Pfenninger**: Writing – review & editing, Writing – original draft, Visualization, Validation, Supervision, Funding acquisition, Resources, Project administration, Methodology, Formal analysis, Data curation, Conceptualization.

## Acknowledgments

The study received support from the LOEWE-Centre Translational Biodiversity Genomics (TBG) funded by Hessen State Ministry of Higher Education, Research and the Arts (HMWK). We would like to thank Ahmet Hulusi Büyükakyüz for his help with the experiment.

