## Supplementary material for "Transcriptomics predicts Artificial Light at Night’s (ALAN) impact on fitness: nightly illumination alters gene expression pattern and negatively affects fitness components in the midge *Chironomus riparius* (Diptera:Chironomidae)"

S1: Candidate genes of significantly differentially expressed genes (p<0.05). Genes were sorted by the functional clusters photoreception, circadian rhythm, development and stress.

| gene id | base mean | L2FC | lfcSE | stat | p | padj | annotation |
| --- | --- | --- | --- | --- | --- | --- | --- |
| photoreception | | | | | | | |
| maker-scaff158-snap-gene-0.494 | 47.57 | 4.26 | 0.94 | 4.54 | 0.0000 | 5.26 | calphotin-like |
| epigenetic regulation | | | | | | | |
| maker-scaff6-augustus-gene-0.116 | 2040.61 | -0.38 | 0.08 | -4.70 | 0.0000 | 5.58 | 25S rRNA (cytosine-C(5))-methyltransferase nop2-like |
| maker-scaff106-augustus-gene-0.288 | 233.49 | 0.53 | 0.19 | 2.74 | 0.0062 | 2.21 | histone-lysine N-methyltransferase trithorax isoform X3 |
| augustus_masked-scaff381-processed-gene-0.7 | 467.94 | -0.49 | 0.17 | -2.93 | 0.0033 | 2.48 | putative methyltransferase C9orf114 |
| maker-scaff211-augustus-gene-1.477 | 2319.75 | -0.44 | 0.13 | -3.37 | 0.0008 | 3.12 | ubiquinone biosynthesis O-methyltransferase, mitochondrial |
| maker-scaff211-snap-gene-1.614 | 6732.03 | 0.33 | 0.09 | 3.53 | 0.0004 | 3.39 | LOW QUALITY PROTEIN: histone-lysine N-methyltransferase 2C-like |
| maker-scaff543-augustus-gene-0.127 | 790.09 | -0.42 | 0.10 | -4.03 | 0.0001 | 4.26 | tRNA (adenine(58)-N(1))-methyltransferase non-catalytic subunit TRM6 |
| maker-scaff198-snap-gene-0.364 | 824.57 | -0.39 | 0.09 | -4.26 | 0.0000 | 4.68 | methyltransferase-like protein 13 |
| maker-scaff317-augustus-gene-0.241 | 1181.03 | -0.33 | 0.08 | -4.29 | 0.0000 | 4.74 | probable histone-lysine N-methyltransferase Mes-4 |
| circadian rhythm | | | | | | | |
| maker-scaff673-augustus-gene-2.280 | 242.97 | 0.45 | 0.14 | 3.20 | 0.0014 | 2.86 | LOW QUALITY PROTEIN: histone acetyltransferase p300 |
| maker-scaff203-snap-gene-0.77 | 2405.64 | 0.41 | 0.11 | 3.68 | 0.0002 | 3.64 | LOW QUALITY PROTEIN: histone acetyltransferase KAT6A |
| maker-scaff741-augustus-gene-0.302 | 14569.66 | 0.54 | 0.11 | 5.00 | 0.0000 | 0.00 | E3 ubiquitin-protein ligase HUWE1 |
| maker-scaff115-snap-gene-0.442 | 942.10 | 1.08 | 0.36 | 2.97 | 0.0030 | 0.03 | nuclear hormone receptor FTZ-F1 isoform X8 |
| maker-scaff211-augustus-gene-1.550 | 6097.77 | 0.39 | 0.12 | 3.20 | 0.0014 | 0.02 | nuclear factor interleukin-3-regulated protein isoform X2 |
| maker-scaff154-augustus-gene-0.437 | 305.65 | 0.39 | 0.25 | -4.20 | 0.0000 | 0.00 | circadian clock-controlled protein daywake-like |
| development | | | | | | | |
| maker-scaff330-augustus-gene-0.97 | 1152.71 | 0.39 | 0.30 | -4.09 | 0.0000 | 0.00 | juvenile hormone acid O-methyltransferase |
| maker-scaff366-augustus-gene-1.213 | 7154.02 | -0.93 | 0.28 | -3.25 | 0.0011 | 0.02 | juvenile hormone epoxide hydrolase 1 |
| snap_masked-scaff9-processed-gene-0.168 | 538.08 | 0.54 | 0.15 | 3.57 | 0.0004 | 0.01 | PREDICTED: ecdysone-induced protein 74EF-like |
| maker-scaff9-augustus-gene-0.236 | 331.15 | 0.67 | 0.18 | 3.66 | 0.0002 | 0.01 | ecdysone-induced protein 74EF isoform X1 |
| maker-scaff634-augustus-gene-0.224 | 1097.99 | -0.69 | 0.25 | -2.76 | 0.0058 | 0.05 | ecdysone 2-monooxygenase |
| maker-scaff576-augustus-gene-4.307 | 516.39 | 0.91 | 0.29 | 3.10 | 0.0019 | 2.71 | cholesterol 7-desaturase nvd |
| maker-scaff424-augustus-gene-0.200 | 65543.92 | 1.31 | 0.48 | 2.74 | 0.0062 | 2.21 | pupal cuticle protein 20-like |
| maker-scaff340-augustus-gene-0.242 | 1202.95 | 1.43 | 0.48 | 2.98 | 0.0029 | 2.54 | pupal cuticle protein 27-like |
| maker-scaff41-augustus-gene-0.312 | 2054.11 | -0.59 | 0.20 | -2.94 | 0.0033 | 0.03 | probable chitinase 1 |
| maker-scaff500-snap-gene-0.474 | 4368.00 | -0.78 | 0.26 | -3.04 | 0.0023 | 0.03 | probable chitinase 1 |
| maker-scaff41-augustus-gene-0.320 | 5749.07 | -0.86 | 0.29 | -2.91 | 0.0036 | 0.03 | probable chitinase 1 |
| maker-scaff35-augustus-gene-0.252 | 5770.05 | 0.95 | 0.30 | 3.23 | 0.0012 | 0.02 | chitin synthase chs-2 isoform X3 |
| maker-scaff317-augustus-gene-0.256 | 20321.42 | 0.98 | 0.35 | 2.77 | 0.0056 | 0.04 | chitin deacetylase 1 |
| maker-scaff422-augustus-gene-0.161 | 205.95 | -0.99 | 0.36 | -2.78 | 0.0055 | 0.04 | probable chitinase 1 |
| maker-scaff86-augustus-gene-0.228 | 2287.01 | -0.81 | 0.28 | -2.84 | 0.0045 | 2.35 | cytochrome P450 6d3-like |
| maker-scaff673-snap-gene-0.318 | 766.30 | -1.15 | 0.39 | -2.97 | 0.0030 | 2.53 | probable cytochrome P450 6a14 |
| maker-scaff576-snap-gene-4.354 | 1296.14 | -0.82 | 0.27 | -3.04 | 0.0024 | 2.62 | probable cytochrome P450 6a14 |
| maker-scaff35-snap-gene-0.301 | 5340.50 | -0.85 | 0.27 | -3.20 | 0.0014 | 2.86 | cytochrome P450 10-like |
| maker-scaff222-augustus-gene-0.400 | 615.48 | -1.11 | 0.34 | -3.32 | 0.0009 | 3.04 | probable cytochrome P450 9f2 |
| maker-scaff17-augustus-gene-0.348 | 262.09 | -0.86 | 0.25 | -3.42 | 0.0006 | 3.21 | cytochrome P450 4c21 |
| maker-scaff222-augustus-gene-0.399 | 817.82 | -0.95 | 0.27 | -3.47 | 0.0005 | 3.28 | probable cytochrome P450 9f2 |
| maker-scaff227-snap-gene-0.177 | 6312.00 | -0.63 | 0.18 | -3.53 | 0.0004 | 3.38 | probable cytochrome P450 28a5 |
| maker-scaff689-snap-gene-0.561 | 282.46 | -0.85 | 0.24 | -3.57 | 0.0004 | 3.45 | probable cytochrome P450 28d1 |
| maker-scaff671-augustus-gene-0.333 | 1379.45 | 0.69 | 0.18 | 3.75 | 0.0002 | 3.75 | cytochrome P450 4c21-like |
| maker-scaff15-augustus-gene-0.273 | 5752.36 | -0.81 | 0.29 | -2.78 | 0.0054 | 2.27 | V-type proton ATPase subunit G |
| maker-scaff628-augustus-gene-0.68 | 1252.65 | 0.53 | 0.18 | 2.95 | 0.0032 | 2.50 | probable phospholipid-transporting ATPase VD isoform X2 |
| maker-scaff33-augustus-gene-1.256 | 1451.25 | -0.20 | 0.06 | -3.26 | 0.0011 | 2.96 | PREDICTED: ATPase family AAA domain-containing protein 1-B |
| maker-scaff643-augustus-gene-0.26 | 2451.96 | 0.52 | 0.14 | 3.72 | 0.0002 | 3.70 | sodium/potassium-transporting ATPase subunit beta-2 isoform X2 |
| maker-scaff117-augustus-gene-0.160 | 17548.48 | 0.41 | 0.11 | 3.80 | 0.0001 | 3.84 | sodium/potassium-transporting ATPase subunit alpha isoform X9 |
| maker-scaff18-snap-gene-0.499 | 4460.73 | 0.36 | 0.09 | 3.85 | 0.0001 | 3.94 | probable phospholipid-transporting ATPase IA isoform X7 |
| maker-scaff154-augustus-gene-0.452 | 2500.80 | 0.38 | 0.09 | 4.08 | 0.0000 | 4.34 | retinal-specific phospholipid-transporting ATPase ABCA4 |
| maker-scaff33-augustus-gene-0.340 | 6569.06 | 0.36 | 0.08 | 4.60 | 0.0000 | 5.38 | plasma membrane calcium-transporting ATPase 2 isoform X1 |
| maker-scaff557-augustus-gene-0.432 | 1116.88 | -0.30 | 0.11 | -2.78 | 0.0054 | 2.26 | probable ATP-dependent RNA helicase DDX56 |
| maker-scaff324-augustus-gene-0.208 | 760.20 | -0.30 | 0.10 | -3.06 | 0.0022 | 2.65 | probable ATP-dependent RNA helicase DDX52 |
| maker-scaff673-augustus-gene-0.237 | 843.86 | -0.22 | 0.07 | -3.14 | 0.0017 | 2.78 | probable ATP-dependent RNA helicase DDX55 homolog |
| maker-scaff175-augustus-gene-0.42 | 1766.76 | -0.28 | 0.08 | -3.61 | 0.0003 | 3.51 | LOW QUALITY PROTEIN: probable ATP-dependent RNA helicase DDX27 |
| maker-scaff364-augustus-gene-0.177 | 821.25 | -0.38 | 0.09 | -4.01 | 0.0001 | 4.22 | ATP-dependent RNA helicase DDX24 |
| maker-scaff734-augustus-gene-0.194 | 1550.55 | -0.46 | 0.11 | -4.27 | 0.0000 | 4.71 | probable ATP-dependent RNA helicase DDX47 |
| stress | | | | | | | |
| maker-scaff645-augustus-gene-0.66 | 179.53 | -1.23 | 0.21 | -5.91 | 0.0000 | 0.00 | glutathione S-transferase 1-1-like |
| maker-scaff607-augustus-gene-0.295 | 1324.01 | -0.64 | 0.19 | -3.46 | 0.0005 | 0.01 | glutathione S-transferase 1-1-like |
| maker-scaff25-augustus-gene-1.508 | 7076.39 | -0.79 | 0.24 | -3.26 | 0.0011 | 0.02 | microsomal glutathione S-transferase 1-like isoform X2 |
| maker-scaff407-augustus-gene-0.6 | 5843.98 | -0.82 | 0.27 | -3.08 | 0.0020 | 0.02 | glutathione S-transferase D7 |
| maker-scaff205-augustus-gene-0.294 | 891.20 | -0.85 | 0.24 | -3.47 | 0.0005 | 0.01 | glutathione S-transferase 1 isoform X1 |
| maker-scaff205-augustus-gene-0.317 | 253.13 | -0.90 | 0.15 | -5.88 | 0.0000 | 0.00 | glutathione S-transferase 1-1 |
| maker-scaff132-snap-gene-0.227 | 65151.86 | -0.76 | 0.20 | -3.85 | 0.0001 | 0.00 | catalase isoform X2 |
| maker-scaff168-augustus-gene-0.410 | 13282.92 | 1.08 | 0.33 | 3.27 | 0.0011 | 2.97 | peroxidase-like isoform X4 |
| maker-scaff31-snap-gene-0.522 | 170.91 | -1.68 | 0.50 | -3.37 | 0.0007 | 0.01 | heat shock 7 kDa protein cognate 2 |
| maker-scaff241-augustus-gene-0.206 | 109112.17 | -0.24 | 0.07 | -3.29 | 0.0010 | 0.01 | heat shock 7 kDa protein cognate 4 |
| maker-scaff378-augustus-gene-0.423 | 6539.93 | -0.47 | 0.12 | -3.99 | 0.0001 | 0.00 | heat shock protein 27-like |
| maker-scaff86-augustus-gene-1.276 | 3567.00 | -0.20 | 0.07 | -2.84 | 0.0045 | 0.04 | activator of 9 kDa heat shock protein ATPase homolog 1 |

S2: Feeding schedule (Doria *et al.* 2021, modified) with amount of food per larvae on the left and total amount of food for 30 larvae suspended in 50ml of

deionised water. Each replicate bowl received 1ml of the suspension daily.

| Day | mg/larvae | Food in 50ml; for 30 larvae |
| --- | --- | --- |
| 1 | 0.1 | 150 |
| 2 | 0.1 | 150 |
| 3 | 0.17 | 255 |
| 4 | 0.17 | 255 |
| 5 | 0.23 | 345 |
| 6 | 0.23 | 345 |
| 7 | 0.3 | 450 |
| 8 | 0.3 | 450 |
| 9 | 0.37 | 555 |
| 10 | 0.37 | 555 |
| 11 | 0.43 | 645 |
| 12 | 0.43 | 645 |
| 13 | 0.5 | 750 |
| 14 | 0.5 | 750 |
| 15 | 0.5 | 750 |
| 16 | 0.5 | 750 |
| 17 | 0.5 | 750 |
| 18 | 0.5 | 750 |
| 19 | 0.5 | 750 |
| 20 | 0.5 | 750 |
| 21 | 0.5 | 750 |
| 22 | 0.5 | 750 |
| 23 | 0.5 | 750 |
| 24 | 0.5 | 750 |
| 25 | 0.5 | 750 |
| 26 | 0.5 | 750 |
| 27 | 0.5 | 750 |
| 28 | 0.5 | 750 |
